# Endogenous BioID elucidates TCF7L1 interactome modulation upon GSK-3 inhibition in mouse ESCs

**DOI:** 10.1101/431023

**Authors:** Steven Moreira, Caleb Seo, Victor Gordon, Sansi Xing, Ruilin Wu, Enio Polena, Vincent Fung, Deborah Ng, Cassandra J Wong, Brett Larsen, Brian Raught, Anne-Claude Gingras, Yu Lu, Bradley W. Doble^✉^

**Affiliations:** Department of Biochemistry and Biomedical Sciences, Stem Cell and Cancer Research Institute, McMaster University, Hamilton, ON L8N 3Z5, Canada; Princess Margaret Cancer Centre, University Health Network, 101 College Street, Toronto, ON M5G 1L7, Canada; Department of Medical Biophysics, University of Toronto, Toronto, ON M5G 1L7, Canada; Lunenfeld-Tanenbaum Research Institute, Mount Sinai Hospital, 600 University Avenue, Toronto, ON M5G 1X5, Canada; Department of Molecular Genetics, University of Toronto, Toronto, ON M5S 1A8, Canada

**Keywords:** BioID, GSK-3, TCF7L1, mESC

## Abstract

Modulation of Wnt target gene expression via the TCF/LEFs remains poorly understood. We employ proximity-based biotin labeling (BioID) to examine GSK-3 inhibitor effects on the TCF7L1 interactome in mouse ESCs. We generated ESC lines with biotin ligase BirA* fused to TCF7L1 by knocking it into the endogenous *TCF7L1* locus or by inserting a doxinducible BirA*-TCF7L1 transgene into the *Rosa26* locus. Induction yielded BirA*-TCF7L1 levels 3-fold higher than in the endogenous system, but substantial overlap in biotinylated proteins with high peptide counts were detected by each method. Known TCF7L1 interactors TLE3/4 and β-catenin, and numerous proteins not previously associated with TCF7L1, were identified in both systems. Despite reduced BirA*-TCF7L1 levels, the number of hits identified with both BioID approaches increased after GSK-3 inhibition. We elucidate the network of TCF7L1 proximal proteins regulated by GSK-3 inhibition, validate the utility of endogenous BioID, and provide mechanistic insights into TCF7L1 target gene regulation.

**Highlights:**

- BirA*-TCF7L1 at single-copy physiological levels generates robust BioID data
- CHIR99021 reduces TCF7L1 levels but increases detectable TCF7L1-proximal proteins
- The TCF7L1 interactome of largely epigenetic/transcription factors fluctuates with GSK-3 inhibition
- JMJD1C, SALL4 and BRG1/SMARCA4 are validated as TCF7L-interacting proteins

## Introduction

Wnt/β-catenin signaling regulates tissue homeostasis and mammalian development. Inappropriate Wnt pathway activation leads to a variety of cancers, most prominently colorectal cancer (1, 2). Wnt ligands and cognate cell surface receptors initiate a cascade of intracellular signaling events that ultimately lead to activation of Wnt target genes via β-catenin interactions with the T-Cell Factor/Lymphoid Enhancer Factor (TCF/LEF) family of transcription factors (3).

In mouse embryonic stem cells (mESCs) all four TCF/LEF family members, TCF7, TCF7L1, TCF7L2, and LEF1 are expressed at detectable protein levels (4, 5), although TCF7L1 is the most abundant and best characterized TCF/LEF in this cell type (6). TCF7L1 is part of an extended network of transcription factors regulating pluripotency (7, 8). Derepression of TCF7L1 is necessary for the enhancement of self-renewal in mESCs in response to Wnt activation (9, 10). Stimulation with the GSK-3 inhibitor CHIR99021 (CHIR) or Wnt3a results in the reduction of TCF7L1 transcript levels as well as β-catenin-dependent proteasomal degradation of TCF7L1 (11).

Despite being extensively studied, the precise mechanisms through which TCF7L1 regulates mESC pluripotency remain poorly characterized. Additionally, few TCF/LEF protein-protein interactions have been described in the literature. In the absence of β-catenin, TCF7L1 is thought to act as a constitutive transcriptional repressor in association with corepressors, including members of the Transducin-Like Enhancer of Split (TLE) family (Groucho in Drosophila) and C-terminal binding protein (CtBP) (6, 10, 12–17). In the presence of a Wnt signal, the TCF/LEFs interact with β-catenin, which recruits co-activators such as P300, CBP, SMARCA4 and BCL9 (18–22).

Many protein-protein interactions have been identified by using affinity purification coupled with mass spectrometry, however, due to the insolubility of chromatin-associated proteins, this technique is restricted to strong interactions and is prone to false positives. Proximity-dependent biotinylation, BioID, is a novel technique for identifying protein-protein interactions in cells (23). This technique employs a promiscuous biotin ligase mutant, BirA*, which is coupled to a “bait” protein allowing for the labelling of vicinal proteins within a radius of approximately 10nm (23, 24). The BioID method allows for vigorous cell lysis, as biotinylated proteins are affinity purified via high-affinity interactions with immobilized streptavidin. The technique also permits detection of transient and/or weak interactions.

Here, we use BioID to identify novel TCF7L1-interacting proteins in differentiating mouse embryonic stem cells, both in the absence or presence of the Wnt mimetic small molecule CHIR. BioID conventionally employs stable cell lines with constitutively or inducibly overexpressed baits. To minimize potential overexpression artefacts, we used homologous recombination to introduce BirA* at the N-terminus of endogenously expressed TCF7L1 and compared our BioID results from this approach with a doxycycline-inducible BirA*-TCF7L1 overexpression system. Our data reveal 146 putative TCF7L1 proximal proteins, offering new insights into mechanisms of TCF7L1 function and the influence of Wnt signaling on the TCF7L1 interactome.

## Results

### A. Inducible and endogenous BirA*-TCF7L1 BioID systems

To ensure a high degree of confidence in our TCF7L1 BioID screen, we first identified the condition that provided us with maximal levels of TCF7L1 protein in wild-type (WT) E14TG2a mouse ESCs. The expression of TCF7L1 was assessed in WT cells cultured in standard medium containing 15%serum plus LIF, medium lacking LIF, and medium with 5% serum and lacking LIF (Figure 1A). The levels of TCF7L1 increased similarly with LIF withdrawal in both serum conditions (Figure 1A), thus we used 5% serum in subsequent experiments to minimize serum effects.

To generate mESCs with inducible BirA*-TCF7L1, we sub-cloned a doxycycline (dox) inducible expression system into a targeting vector directing homologous recombination within the safe harbour *Rosa26* locus. We employed a TALEN-facilitated knock-in approach to introduce a single copy of dox-inducible 3xFLAG-BirA*-P2A- or 3xFLAG-BirA*-tagged TCF7L1 into the *Rosa26* locus. We will refer to the TCF7L1 inducible lines as BirA*-P2A-TCF7L1 RT, a control cell line incorporating a P2A self-cleaving peptide between the BirA* and TCF7L1 coding sequences, or BirA*-TCF7L1 RT (the RT suffix denotes Rosa TetOne).

Expression of inducible proteins in RT clones was confirmed by western blotting (Figs. 1B, 1D, S1A). Addition of 1µg/mL doxycycline induced expression of both the BirA*-P2ATCF7L1 RT control and the BirA*-TCF7L1 RT fusion protein (Figs. 1B, S1A). Treatment with CHIR reduced TCF7L1 protein levels significantly (Figure 1B). Total TCF7L1 levels were elevated in BirA*-TCF7L1 RT lines after doxycycline addition, whereas dox had no effect on TCF7L1 levels in wild-type mESCs.

Unexpectedly, after dox treatment BirA*-P2A-TCF7L1 RT lines also demonstrated more detectable TCF7L1 at 70 kDa. BirA*-P2A-TCF7L1 RT controls also displayed a detectable uncleaved BirA*-P2A-TCF7L1 fusion product of approximately 130 kDa (upper panel, Figure 1B). The levels of BirA* (40 kDa) in the BirA*-P2A-TCF7L1 RT line and BirA*-TCF7L1 fusion protein (130 kDa) in the BirA*-TCF7L1 RT line were reduced in response to CHIR, suggesting post-transcriptional regulation of TCF7L1 (Figure 1B). To generate the endogenous TCF7L1 BioID system, we used TALEN-facilitated homologous recombination to introduce a single copy of 3xFLAG-BirA*-P2A- and 3xFLAG-BirA*-tagged TCF7L1 at the endogenous *TCF7L1* locus. We will refer to the TCF7L1 endogenous lines as BirA*-P2ATCF7L1 or BirA*-TCF7L1.

Expression of BirA*-P2A-TCF7L1 or BirA*-TCF7L1 in targeted clones was confirmed by western blotting (Figs. 1C, S1B). We did not observe appreciable changes in total TCF7L1 protein levels in either BirA*-P2A or BirA*-TCF7L1 mESC lines. A reduction in TCF7L1 was observed in the presence of CHIR (Figure 1C). Again, we observed a minimal amount of uncleaved BirA*-P2A-TCF7L1 in BirA*-P2A controls. As observed in the dox-inducible system, BirA* levels in control cell lines were reduced in the presence of CHIR in the endogenous system.

Expression levels of BirA* fusion proteins in the two BioID systems were compared by western blotting (Figure 1D). The inducible system demonstrated a significant increase in both BirA*-P2A-TCF7L1 and BirA*-TCF7L1 protein levels compared to the endogenous system (Figure 1D).

### B. Inducible and endogenous TCF7L1 BioID systems demonstrate biotinylation of proteins in vivo

We observed heterogeneous expression and nuclear localization of BirA* in BirA*-P2A-TCF7L1 RT and BirA*-TCF7L1 RT mESC lines when anti-FLAG immunofluorescent staining was visualized (Figure 2A). However, BirA*-P2A-TCF7L1 RT cells, in addition to nuclear fluorescence, displayed diffuse cytosolic signal. Although treatment with CHIR caused a reduction in total TCF7L1 protein (Figure 1B), there were many individual cells that retained elevated levels of TCF7L1 (Figure 2A).

We used fluorescently conjugated streptavidin to detect levels of biotinylation in WT, BirA*-P2A-TCF7L RT and BirA*-TCF7L1 RT mESC lines induced with doxycycline in the absence or presence of biotin and/or CHIR (Figure 2A). Streptavidin staining revealed a considerable amount of background biotinylation in wildtype mESCs, even in the absence of biotin. Despite appreciable levels of background biotinylation, we observed elevated biotinylation in BirA*-P2A-TCF7L1 RT and BirA*-TCF7L1 RT mESCs with the highest levels of detectable BirA* (Fig 2A).

We used the same immunofluorescence approach to examine the biotinylation activity in endogenous BirA* P2A-TCF7L1 and BirA*-TCF7L1 mESC lines (Figure 2B). BirA* expression as assessed by FLAG immunofluorescence, was lower and more homogeneous in the endogenous system when compared to the inducible system. As in the inducible system, BirA* P2A-TCF7L1 and BirA*-TCF7L1 localized to the nucleus, with the P2A controls also demonstrating diffuse cytosolic staining. Treating with CHIR led to an overall reduction in TCF7L1, however, again we observed individual cells where TCF7L1 levels remained elevated. In contrast to the inducible BioID system, we did not observe any visible increase in biotinylation in BirA*-P2A or BirA*-TCF7L1 cell lines, as assessed by streptavidin staining.

To detect biotinylation activity with higher sensitivity, we used chemiluminescent visualization of streptavidin-HRP. In the inducible BioID system we observed biotinylation of unidentified proteins as well as auto-biotinylation of BirA*-P2A-TCF7L1 RT and BirA*-TCF7L1 RT fusions in the absence and presence of CHIR (Figure 2C). Despite an undetectable increase in biotinylation as assessed by immunofluorescence, we observed auto-biotinylation of BirA*-P2ATCF7L1 and BirA*-TCF7L1, as well as biotinylation of unidentified proteins, in the endogenous BioID system (Figure 2D).

**Fig. 1.**
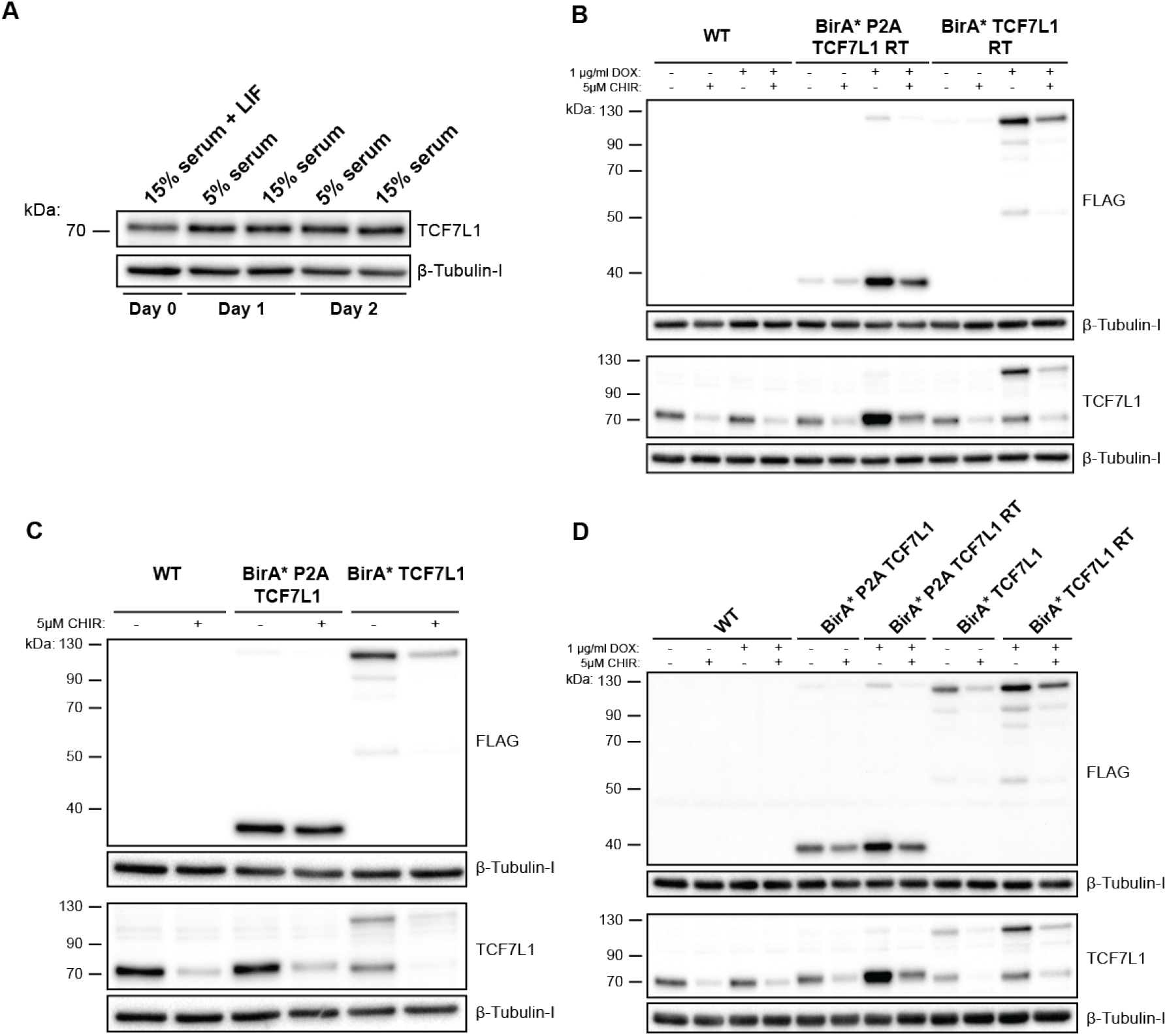
BirA*-P2A-TCF7L1 and BirA*-TCF7L1 expression in inducible and endogenous BioID systems. (A) Western blot analyses of WT mESCs maintained for up to 48h in 5% or 15% serum, as indicated. Lysates were probed with antibodies against TCF7L1 and β-Tubulin-I, as indicated. (B) Western blots obtained with lysates from BirA*-P2A-TCF7L1 RT, BirA*-TCF7L1 RT and WT cells maintained in 5% serum for 24h and subsequently treated with 5µM CHIR and/or 1µg/mL DOX for 24h. Lysates were probed with antibodies against FLAG, TCF7L1, and β-Tubulin-I, as indicated. (C) Western blots assessing lysates from BirA*-P2A-TCF7L1, BirA*-TCF7L1 and WT cells maintained in 5% serum for 24hr and then treated with 5µM CHIR, for 24h. Lysates were probed with antibodies against FLAG, TCF7L1, and β-Tubulin-I, as indicated. (D) Western blots obtained with lysates from WT, BirA*-P2A-TCF7L1, BirA*-P2A-TCF7L1 RT, BirA*-TCF7L1, and BirA*-TCF7L1 RT cells maintained in 5% serum for 24hr and then treated with 5µM CHIR and/or 1µg/mL DOX, for 24h. Lysates were probed with antibodies against FLAG, TCF7L1, and β-Tubulin-I, as indicated.

### C. BioID-based assembly of the TCF7L1 interactome

As we observed biotinylation of unidentified proteins in the inducible and endogenous BioID systems, we were confident in our ability to identify these proteins in a large-scale BioID pull-down followed by mass-spectrometry. Cells were cultured in 5% serum for an initial 24 hours, followed by supplementation with biotin, with or without CHIR for 24 hours. For the inducible system, cells were also treated with dox at the time of biotin addition. BioID was performed on 3 biologically independent clones for each expression system including the BirA*-P2A-TCF7L1 controls. We isolated biotinylated proteins via streptavidin-biotin affinity purification, and subsequently analyzed them by using LC-MS/MS mass spectrometry. Data were searched using the ProHits suite, and SAINTexpress was used to calculate the probability of each potential proximal-protein relative to control samples.

Both BioID screens identified Wnt pathway components. Known TCF7L1-associated co-repressor proteins TLE3 and TLE4 were among the biotinylated proteins detected, in the absence and presence of CHIR (Figure 3A, 3B). In the endogenous BioID screen both proteins were of lower confidence in the absence of CHIR and more abundant, with a higher confidence, in its presence (Figure 3A, 3B). By contrast, in the inducible BioID screen, we observed an enrichment of both TLE3 and TLE4 in the absence of CHIR and lower levels in its presence (Figure 3A, 3B). Despite their acknowledged role in TCF/LEF-mediated control of transcription, TLE3 and TLE4 were not highly abundant in either BioID screen.

**Fig. 2.**
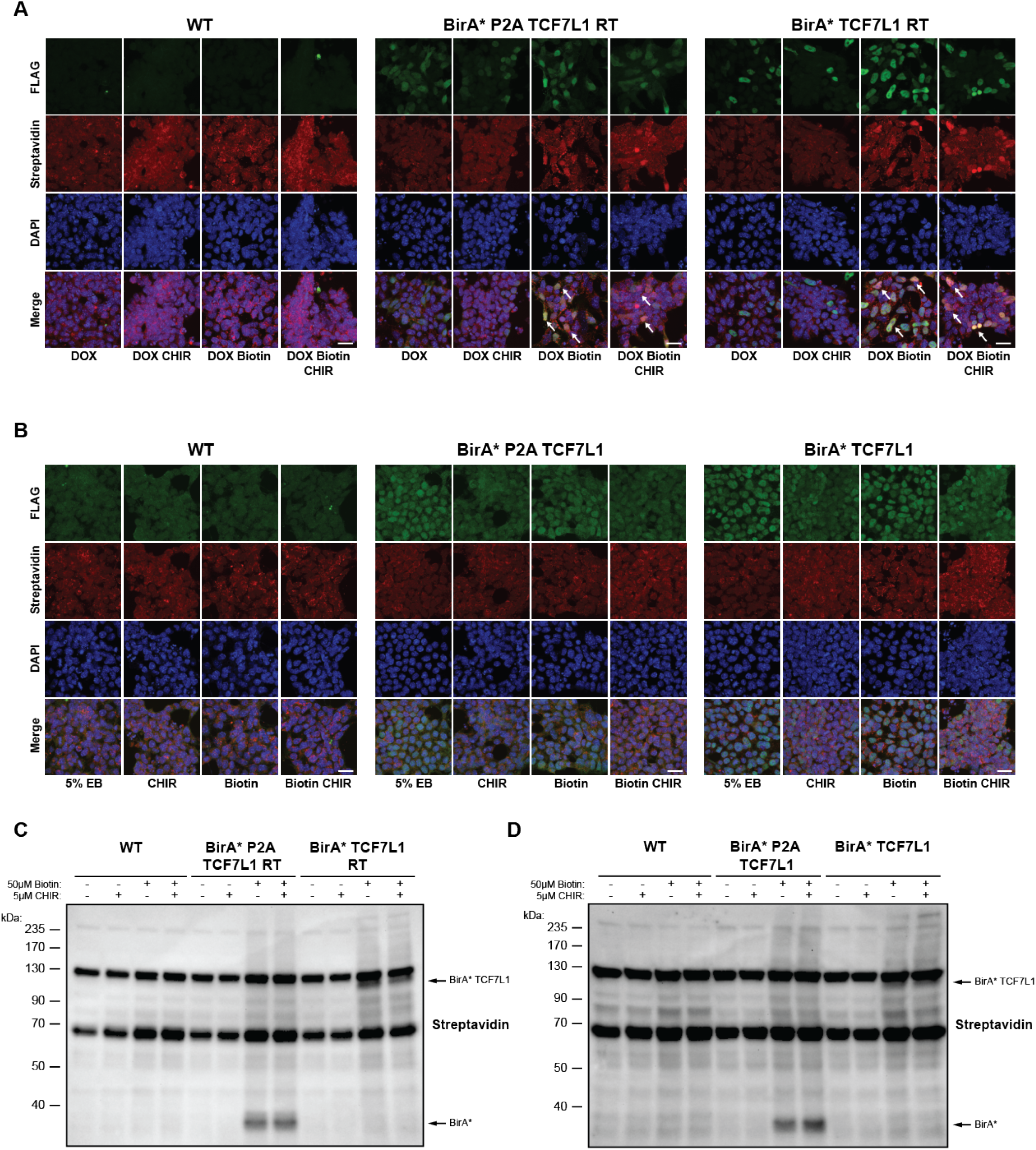
Inducible and endogenous TCF7L1 BioID systems demonstrate biotinylation of proteins in vivo. (A) immunofluorescence images of BirA*-P2A-TCF7L1 RT, BirA*-TCF7L1 RT and WT mESCs maintained in 5% serum for 24h followed by supplementation with combinations of 5µM CHIR, 1µg/mL doxycycline, and 50µM biotin, for 24h. Staining was performed with anti-FLAG (green), fluorophore-conjugated streptavidin (red) and DAPI. Scale bar, 20µm. White arrowheads mark cells with elevated levels of FLAG-tagged BirA* or FLAG-tagged BirA*-TCF7L1 as well as high levels of detectable biotinylation. (B) immunofluorescence images of BirA*-P2A-TCF7L1, BirA*-TCF7L1 and WT cells maintained in 5% serum for 24h, and then treated with 5µM CHIR and/or 50µM biotin, for 24h. Staining was performed with anti-FLAG (green), fluorophore-conjugated streptavidin (red) and DAPI. Scale bar, 20µm. (C) Western blot analysis of biotinylated proteins in lysates of BirA*-P2A-TCF7L1 RT, BirA*-TCF7L1 RT or WT cells maintained in 5% serum for 24h and then treated with 5µM CHIR and/or 50µM biotin, in the presence of 1µg/mL doxycycline, for 24h. Lysates were probed with streptavidin-HRP. Arrows point to 3xFLAG-BirA* fusion proteins. (D) Western blot analysis of biotinylated proteins in a single clone of BirA*-P2A-TCF7L1, BirA*-TCF7L1 and WT cells maintained in 5% serum for 24h, followed by supplementation with 5µM CHIR or 50µM biotin, for 24h. Lysates were probed with streptavidin-HRP. Arrows point to 3xFLAG-BirA* fusion proteins.

The most abundant Wnt signaling component detected in the screens was β-catenin, identified in the presence of CHIR in both systems (Figure 3A, 3B). The inducible BioID screen also detected additional Wnt-associated proteins, including BCL9, CDH1, P300, and BCL9L, which were more abundant in the presence of CHIR but also detected in its absence (Figure 3B). APC and CBP were enriched exclusively in the presence of CHIR (Figure 3B).

In both BioID approaches, CHIR treatment resulted in identification of a higher number of TCF7L1 proximal proteins with elevated peptide counts (Figure 3A, 3B). This is despite lowered TCF7L1 ‘bait’ protein levels (and detected peptides) in the presence of CHIR.

In addition to Wnt-associated proteins, we identified many complexes involved in the regulation of transcription, including the BAF chromatin remodeling complex, which was the most abundant complex identified in both BioID screens (Figure 3A, 3B). BAF components ARID1A, SMARCA4, and SMARCC1, were detected in both BioID screens, with and without CHIR treatment but were enriched in the CHIR condition (Figure 3A, 3B). Another multi-subunit complex we observed in both BioID screens was the nuclear receptor corepressor complex (Figure 3A, 3B). NCOR1, NCOR2, TBL1X, TBL1XR1, and SPEN were enriched in the presence of CHIR (Figure 3A, 3B). In the absence of CHIR, NCOR1 was detected in both BioID screens albeit at much lower levels (Figure 3A, 3B). We also identified members and putative members of the polycomb group proteins, such as BRCA associated protein-1 (BAP1), BCL-6 Corepressor (BCOR), ASXL Transcriptional Regulator 2 (ASXL2) and Lysine demethylase 1B (KDM1B); as well as components of the mixed lineage leukemia (MLL) complex in both BioID screens (Figure 3A, 3B).

Along with complexes involved in transcriptional regulation we also identified numerous transcriptional regulators, including JMJD1C, an H3K9me2 demethylase, which was the most abundant hit observed in both screens +/− CHIR (Figure 3A, 3B). Four other abundant transcriptional regulators identified in both BioID screens were, two Spalt transcription factors SALL4 and SALL1; TCF20, a transcriptional coactivator; and ARID3B, a context-dependent transcriptional activator or repressor (Figure 3A, 3B).

**Fig. 3.**
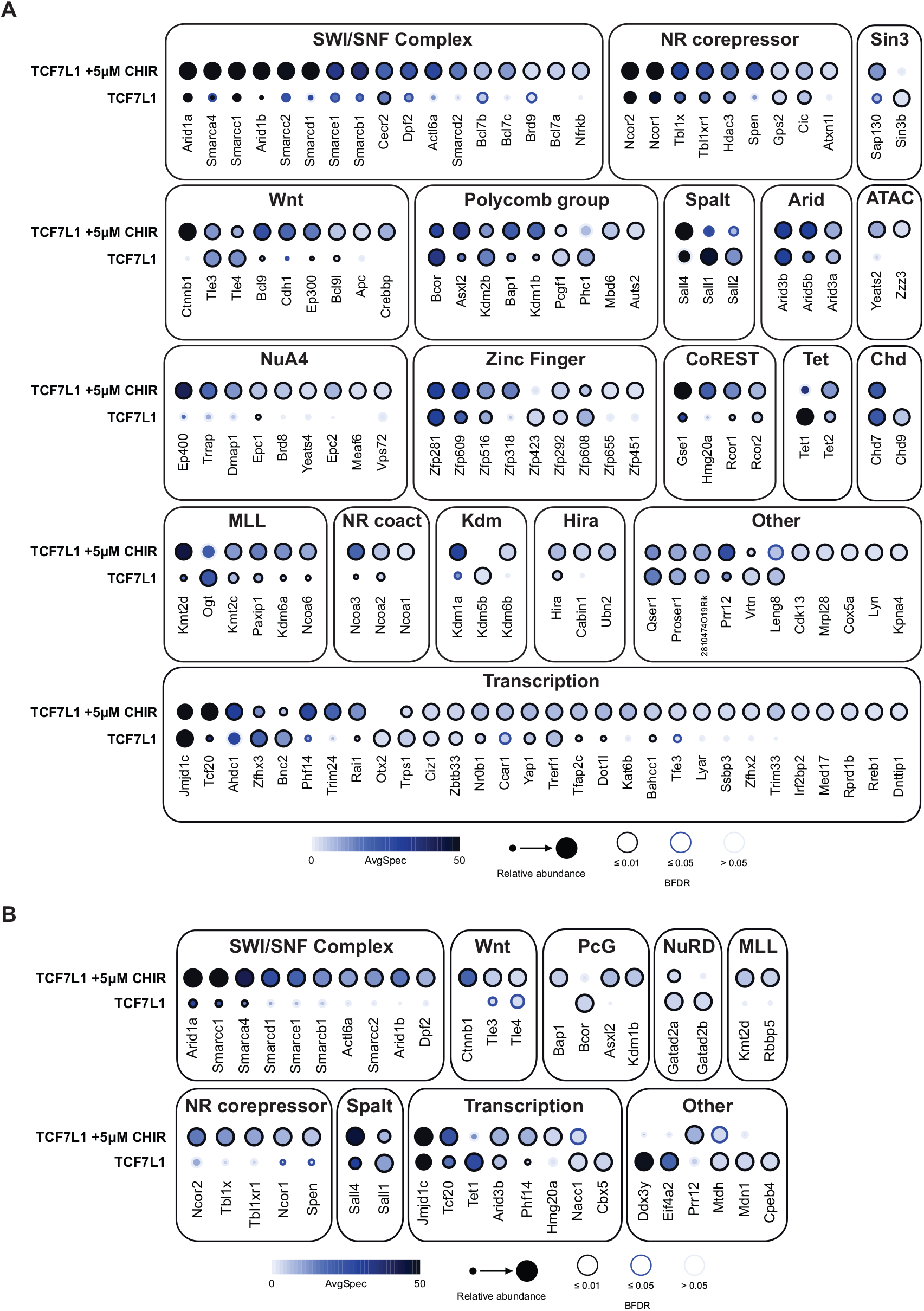
Proteins biotinylated by BirA*-TCF7L1 in inducible and endogenous BioID systems in the presence and absence of CHIR. (A) Dot plot of TCF7L1-proximal proteins identified by using the inducible BioID system in mESCs cultured in 5% serum for 24h and then treated with 50µM biotin, 1µg/mL DOX and 5µM CHIR for 24h. (B) Dot plot of TCF7L1-proximal proteins identified by using the endogenous BioID system in cultured in 5% serum for 24h and then treated with 50µM biotin and 5µM CHIR for 24h. In both systems, all biotinylated proteins identified demonstrated a BFDR ≤ 0.05 and a SAINT score > 0.8 in at least one of the 2 conditions. Spectral counts were capped at 50.

### D. Inducible and endogenous BioID screens detect similar, but distinct, TCF7L1 interactomes

Although our main objective was identifying TCF7L1 proximal and potentially interacting proteins, we also wanted to compare single-copy endogenous BioID with commonly used inducible overexpression of single-copy BioID transgenes. Both approaches were able to identify similar proximal proteins when considering the most abundant statistically significant top hits. Of the 43 proteins identified in the endogenous screen with and without CHIR, only 10 were unique to the endogenous BioID screen, indicating a large overlap in hits detected by the two systems.

The inducible BioID screen was able to identify more low-abundance hits (Figure 3A, 4A). Specifically, in the absence of CHIR, the inducible screen identified 7 unique proteins, of which only OGT, an O-GlcNAc transferase, was of moderate abundance (Figure 4A, 4B). Similarly, in the presence of CHIR, the inducible screen identified 43 unique proteins, which included AHDC1, EP400, SMARCD2, TRIM24, and TRRAP (Figure 4A, 4B). In contrast to the inducible BioID screen, the endogenous BioID screen produced a more tractable list of candidate TCF7L1 proximal proteins (Figure 4A, 4B).

**Fig. 4.**
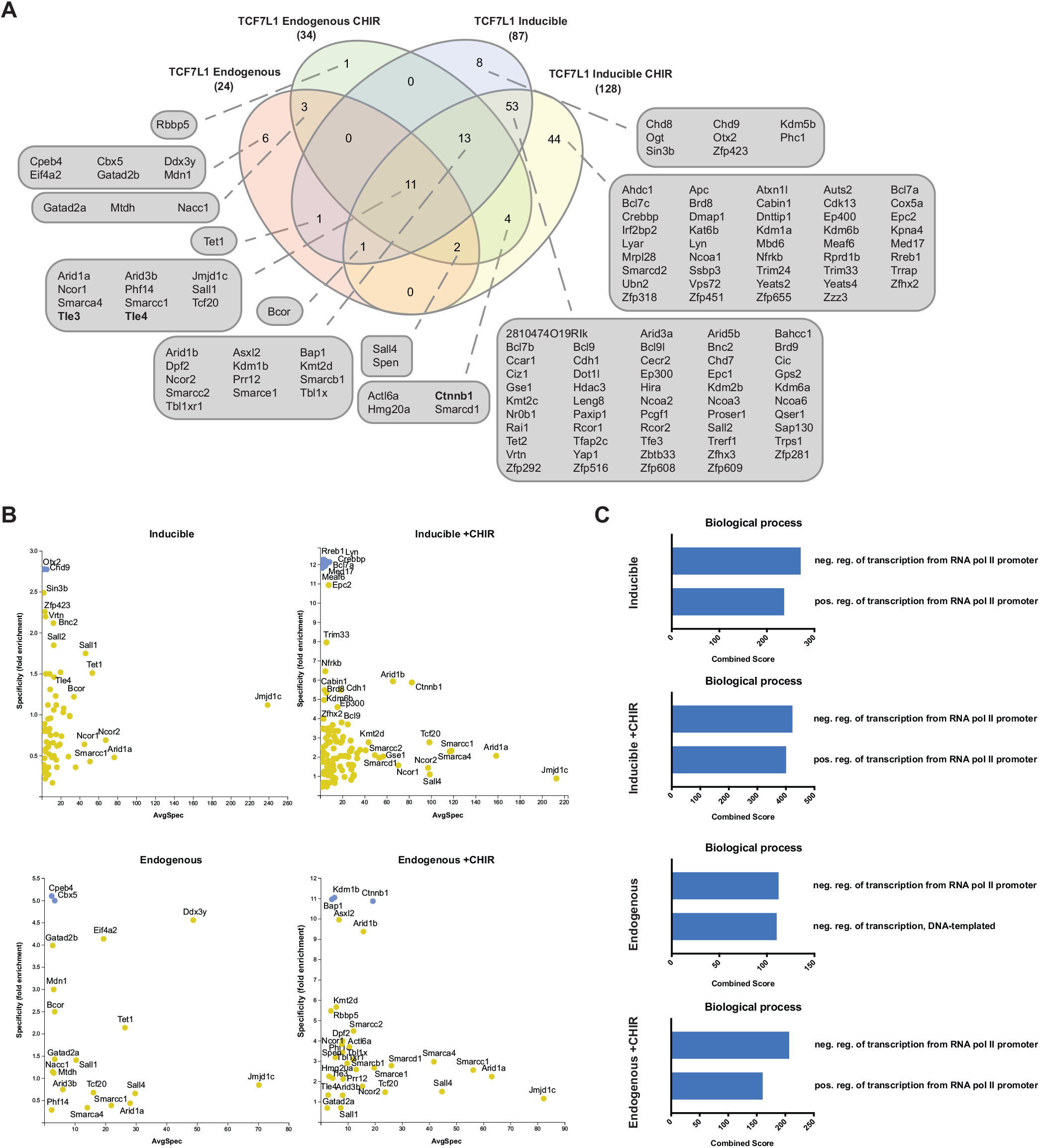
Inducible and endogenous BioID screens detect similar, but distinct, TCF7L1 interactomes. (A) Venn diagram showing overlapping biotinylated proteins identified in the endogenous and inducible BioID systems in the absence or presence of 5µM CHIR, as indicated. Previously known TCF7L1-interacting proteins are highlighted in bold. (B) Prey specificity graphs of TCF7L1-proximal proteins identified by using either the endogenous or inducible BioID systems in the absence or presence of 5µM CHIR, as indicated. Fold enrichment is calculated based on the spectral counts of each prey vs TCF7L1 in each condition, relative to the entire dataset for both conditions. A BDFR cutoff of 0.01 was used. Blue circles indicate preys with infinite specificity (C) Gene ontology of the TCF7L1 proximal proteins identified by using endogenous and inducible BioID systems in absence or presence of 5µM CHIR, as indicated.

In the absence of CHIR, the endogenous screen identified 6 unique proteins, all of low abundance: CPEB4, CBX5, DDX3Y, EIF4A2, GATAD2B, and MDN1. In the presence of CHIR, the endogenous screen identified 1 unique protein, RBBP5 (Figure 4A, 4B).

Although we observed unique proteins in both screens, with or without CHIR, the majority were of low abundance. Focusing on abundant proteins in the absence of CHIR, the only protein observed in both screens was TET1 (Figure 4A, 4B), a methylcytosine dioxygenase involved in DNA demethylation. Conversely, with CHIR treatment we detected β-catenin; two components of the BAF complex, ACTL6A and SMARCD1; and a CoREST subunit, HMG20A; in both screens (Figure 4A, 4B).

Eleven proteins were observed in all conditions in both BioID screens: 3 subunits of the BAF complex ARID1A, SMARCA4, and SMARCC1; TLE3 and TLE4; and ARID3B, JMJD1C, NCOR1, PHF14, SALL1, and TCF20 (Figure 4A, 4B). With the exception of the TLEs, the levels of most of these proteins were unchanged or enriched in the presence of CHIR (Figure 3A, 3B). SALL1 was observed at lower levels in both screens in the presence of CHIR (Figure 3A, 3B).

Thirteen proteins were identified in all screens except the endogenous screen in the absence of CHIR: 5 BAF subunits (ARID1B, DPF2, SMARCB1, SMARCC2, and SMARCE1); 3 polycomb group proteins (ASXL2, BAP1 and KDM1B); 3 components of the nuclear receptor corepressor complex (NCOR2, TBL1X, and TBL1XR1); and KMT2D and PRR12 (Figure 4A, 4B). All of these proteins were detected by the inducible BioID screen without CHIR, albeit at much lower levels.

In total, we identified 146 potential TCF7L1 interacting proteins in the absence and presence of Wnt/β-catenin pathway stimulation via CHIR. Gene ontology (GO) analysis revealed terms associated with negative regulation of transcription in both screens, with and without CHIR (Figure 4C). In the absence of CHIR, the inducible BioID screen contained terms associated with both positive and negative regulation of transcription, in contrast to the endogenous BioID screen, which only contained repressive factors (Figure 4C). GO terms in both BioID screens performed in the presence of CHIR were associated with positive and negative regulation of transcription (Figure 4C).

### E. Validation of JMJD1C, SALL4, and SMARCA4 interactions with TCF7L1 in mESCs

The putative TCF7L1 interactors JMJD1C, SALL4, and SMARCA4 were abundantly identified in the endogenous and inducible BioID screens in the absence or presence of CHIR stimulation. We employed the proximity ligation assay (PLA) to gain additional evidence for the existence of interactions between these proteins and TCF7L1. With this technique, interacting proteins appear as fluorescent foci based on the close proximity of specific antibodies against each of the putative interacting proteins. To facilitate our characterization of putative TCF7L1-interacting proteins, we generated 3xFLAG-TCF7L1 knock-in mESC lines in which the 3xFLAG epitope was added to the N-terminus of one allele of endogenous TCF7L1. FLAG- or JMJD1C- specific antibodies, when used individually in PLA assays using 3xFLAG-TCF7L1 knock-in mESCs, resulted in background levels of fluorescence (Figure 5A). However, when these cells were probed with both FLAG and JMJD1C antibodies, a significant increase in predominantly nuclear punctae was observed (Figure 5A).

**Fig. 5.**
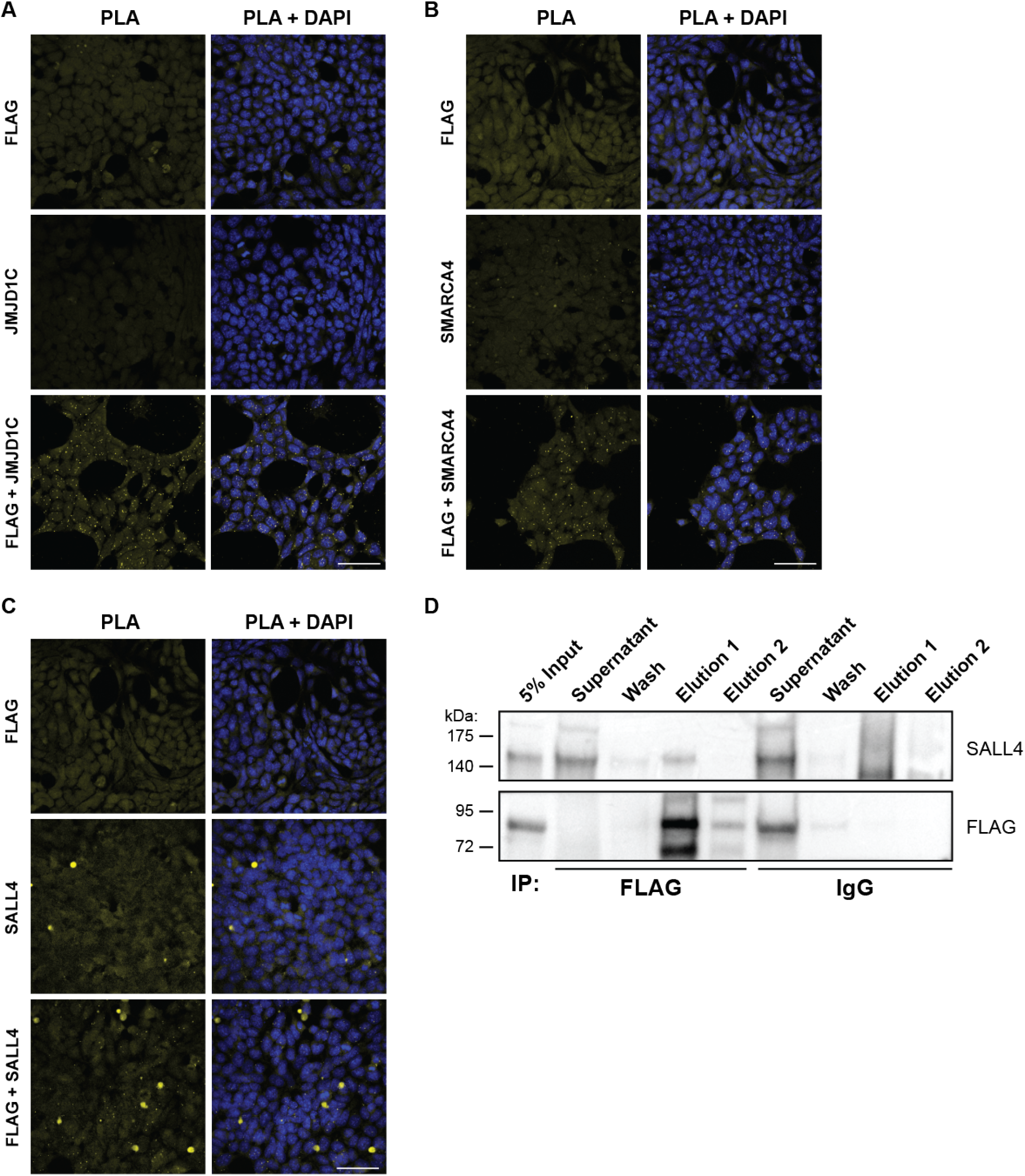
Validation of JMJD1C, SALL4, and SMARCA4 interactions with TCF7L1 in mESCs. (A) PLA results obtained by using 3xFLAG-TCF7L1 knock-in mESCs cultured in standard medium (serum and LIF) and anti-FLAG and anti-JMJD1C primary antibodies or single antibody controls. Nuclei were visualized with DAPI. Scale bar, 50µm. (B) PLA results obtained by using 3xFLAG-TCF7L1 knock-in mESCs cultured in standard medium (serum and LIF) and anti-FLAG and anti-SMARCA4 primary antibodies or single antibody controls. Nuclei were visualized with DAPI. Scale bar, 50µm. C) PLA results obtained by using 3xFLAG-TCF7L1 knock-in mESCs cultured in standard medium (serum and LIF) and anti-FLAG and anti-SALL4 primary antibodies or single antibody controls. Nuclei were visualized with DAPI. Scale bar, 50µm. D) Western blot data for coimmunoprecipitations conducted with anti-FLAG pull-downs and subsequent FLAG and SALL4 immunoblotting using lysates from mESCs with a 3xFLAG epitope knocked into the endogenous TCF7L1 locus.

Similarly, when 3xFLAG-TCF7L1 knock-in mESCs were probed with FLAG- or SMARCA4- specific antibodies individually, background levels of fluorescence with few punctae were detected, whereas numerous strong nuclear punctae were observed when both FLAG and SMARCA4 antibodies were used together (Figure 5B).

We obtained similar PLA results when assaying for interactions between SALL4 and FLAG-tagged TCF7L1, although the number of punctae was fewer and some cells did not display any punctae. (Figure 5C). The observation of fewer punctae is consistent with the BioID data (Figure 3), which suggest less interaction between TCF7L1 and SALL4 in the absence of CHIR, the condition in which the PLA experiments were conducted. Nonetheless, in co-immunoprecipitation experiments, we obtained convincing western blot data suggesting that SALL4(isoform SALL4A, but not SALL4B) and FLAG-tagged TCF7L1 interact (Figure 5D). By contrast, we were unable to obtain co-immunoprecipitates between 3xFLAG-TCF7L1 and JMJD1C or SMARCA4.

## Discussion

Our data suggest that TCF7L1 interacts with numerous transcriptional regulators and chromatin remodeling complexes both in the presence and absence of CHIR-mediated Wnt pathway activation in mESCs. We used mESCs for our study for the following reasons: 1) mESCs express high levels of TCF7L1, 2) Wnt signals regulate TCF7L1 function and stability in mESCs through poorly understood mechanisms, and 3) Identification of putative TCF7L1 interactors by using conventional transgenic as well as endogenous single-copy BioID approaches provides biological insights into mechanisms of TCF7L1 function and highlights the consequences of even small variations in expression of BioID baits. There was significant overlap in the biotinylated proteins identified by using both BioID methods, but the modest 2- to 3-fold increase in BirA*-TCF7L1 in the inducible system yielded approximately one hundred putative TCF7L1 proximal proteins that were undetected with the endogenous system.

TCF7L1 is potent in its ability to promote mESC differentiation, and its levels are strictly controlled (6, 10, 11, 25). It is not surprising that even low-level overexpression yielded different BioID data from the endogenous system and that the expression of more “bait” identified more hits. As the endogenous BioID screen contains physiological levels of TCF7L1, we suggest that its hits are more likely to be biologically relevant. A smaller list of candidate TCF7L1 proximal proteins facilitates subsequent validation and helps build a meaningful interactome with less noise contributed by false-positive hits that are likely in an overexpression system.

Our introduction of BirA* at the N-terminus of endogenous TCF7L1 provides another distinct advantage in that all splice-variant isoforms of TCF7L1 present in mESCs will serve as BioID baits, whereas conventional inducible over-expression systems are restricted to a single isoform.

β-catenin binding to the TCF/LEF factors was once thought to result in the displacement of TLEs, followed by recruitment of co-activators and Wnt target gene activation (19, 21, 26). This model has been challenged as it was demonstrated that β-catenin and TLE1 bind to independent regions and do not compete for binding to TCF7L1 (27). Furthermore, through BioID and co-immunoprecipitation experiments it has been demonstrated that the co-activator BCL9-related β-catenin-binding protein, B9L, is able to form a complex with TCF7L2 together with TLE3 and TLE4, in both the presence and absence of Wnt stimulation in HEK293 cells (28).

Our BioID screens suggest that TCF7L1 associates with TLE3 and TLE4 with and without CHIR, consistent with current working models. Notably, TLE3 and TLE4 were not highly abundant TCF7L1 vicinal proteins in either screen. Intriguingly, the changes observed in TLE3 and TLE4 binding in the absence or presence of CHIR were different between the two BioID systems. Unexpectedly, in the endogenous BioID screen both TLE3 and TLE4 were enriched in the presence of CHIR, despite a downregulation of TCF7L1 protein levels. This is consistent with a B9L BioID screen that identified an increase in TLE3 and TLE4 after stimulation with Wnt-conditioned media (28). By contrast, in the inducible system, TLE3 and TLE4 were reduced with CHIR treatment. This discrepancy could be due to a change in the stoichiometry of Wnt enhanceosome complex components arising from supra-physiological TCF7L1 levels in the inducible cells.

A putative TCF7L1-associated protein that was highly abundant in both screens, specifically in the absence of CHIR, was TET1. TCF7L1 has also been shown to promote the transition from naïve to primed pluripotency (29). TET1 is of particular interest, as together with ZFP281, identified in the inducible BioID screen, it promotes the transition from the naïve to primed pluripotent states (30). Future studies will be needed to determine potential interplay between TET1 and TCF7L1 in the regulation of naïve to primed pluripotency.

The three proteins we chose for validation purposes include the H3K9 demethylase JMJD1C, the chromatin remodeling factor BRG1/SMARCA4, and the transcription factor SALL4. Recently, it has been reported that JMJD1C is required for the maintenance of embryonic stem cell self-renewal and is downregulated during differentiation from the pluripotent state (31). SALL4 is also essential for maintenance of self-renewing pluripotent stem cells and is lost upon ESC differentiation (32–34). Intriguingly, *SALL4* has been reported to be directly activated by TCF/LEF binding to the *SALL4* promoter (35). Conversely, SALL4 is a direct Wnt target gene and both SALL4 isoforms (A and B) have been shown to associate with β-catenin, enhancing TCF reporter activity (35, 36). Our successful co-immunoprecipitation of SALL4A and TCF7L1 suggests that they are not only proximal to each other, they interact with each other, either directly or indirectly, in a complex. Our PLA data also suggest that the same may be true for JMJD1C and TCF7L1.

The most abundant complex identified in both BioID screens was the BAF complex. Components of the BAF complex have been shown to exert differential effects on Wnt target genes. OSA a Drosophila ortholog of ARID1A, was found to be necessary for repression of Wingless (Drosophila ortholog of Wnt) target genes (37). SMARCA4 has been shown to interact with β-catenin, promoting Wnt target gene activation in a TCF-dependent manner (18). Furthermore, recent work demonstrated that SMARCA4 was required for activation of Wnt target genes and deletion of SMARCA4 prevented Wnt-driven tumorigenesis in the small intestine of mice (38). SMARCC1 and multiple BAF subunits, identified as negative regulators of NANOG in an RNAi screen, were shown to promote heterochromatin and chromatin compaction at the NANOG locus (39). Similarly, TCF7L1 has also been shown to bind to the NANOG locus, repressing its expression (6–8). Taken together, our BioID screen suggests that in the absence of a Wnt signal, TCF7L1 recruits BAF complex members at target genes to promote chromatin compaction and subsequent transcriptional repression.

In our BioID screens we observed an enrichment in TCF7L1 association with BAF components during CHIR stimulation, despite post-transcriptional downregulation of TCF7L1 (5, 11). This could be explained by recruitment of additional BAF complex subunits mediated by the TCF7L1-β-catenin interaction. In particular SMARCD1 was only observed in presence of CHIR in both TCF7L1 BioID screens. Notably, it is part of an ESC-specific BAF complex, which facilitates the activation of pluripotency associated genes (40).

TCF7L1 repression of Wnt target genes can be alleviated through β-catenin-dependent TCF7L1 degradation and subsequent derepression of target genes (11, 41). However, the precise mechanism through which β-catenin promotes proteasomal degradation of TCF7L1 remains elusive. A complex identified in our BioID screen that offers potential insights into this mechanism is the nuclear receptor corepressor complex (NCoR). Both TBL1X and TBL1XR1 have been shown to mediate transcriptional activation by recruiting the ubiquitin-conjugating/19S proteasome complex, which enables a switch from corepressor to coactivator usage (42–44). TBL1X has also been linked to proteasomal degradation of β-catenin in response to genotoxic stress-mediated p53 activation, forming a SCF-like E3 complex consisting of SIAH1, SIP, SKP1, APC and TBL1X (Matsuzawa and Reed, 2001). Conversely, TBL1X and TBL1XR1 also interact with β-catenin in response to Wnt signals, binding to target gene promoters to mediate transcriptional activation (45).

Whether TBL1X and TBL1XR1 regulate TCF7L1 protein levels and subsequent Wnt target gene transcription remains to be elucidated. We propose that β-catenin binding to TCF7L1 could mediate the recruitment of TBL1XTBL1XR1, leading to subsequent proteasomal degradation of TCF7L1 and/or its associated corepressors. Subsequently, target genes could either be derepressed or activated via recruitment of co-activators. In support of this TBL1XTBL1XR1 were only associated with TCF7L1 in the presence of CHIR in the endogenous BioID screen. Although TBL1X-TBL1XR1 associated with TCF7L1 without CHIR in the inducible screen, this could be a compensatory mechanism to reduce abnormally elevated levels of TCF7L1 protein.

When determining the effects of the TCF/LEFs on Wnt target genes, the emphasis has been on transcriptional regulation conferred by the corepressor TLEs and co-activators recruited by β-catenin, in the absence and presence of a Wnt signal, respectively. Our TCF7L1 BioID data strongly suggest that TCF7L1 associates more abundantly with numerous other transcriptional regulators and complexes. Furthermore, our BioID screens provide insight into TCF7L1 associated complexes capable of mediating post-transcriptional regulation of TCF7L1 levels, as well as β-catenin-mediated derepression and/or activation of target genes. Future work interrogating the precise mechanisms of TCF7L1 regulation by TET1 and the BAF and NCoR complexes will be instrumental in determining how Wnt-β-catenin signaling mediates derepression/activation of TCF7L1, to regulate stem cell self-renewal and pluripotency.

## Experimental Procedures

Unless otherwise stated, chemicals were obtained from Sigma-Aldrich.

### Proximity Ligation Assay

We used the Duolink^®^ In Situ Red Starter kit (Sigma), with cells grown on ibidi µ-slide 8 well chamber slides (ibiTreat surface), according to manufacturer’s instructions. Pro-Long Gold antifade (containing DAPI) solution (Life Technologies) was used to preserve processed cells (1–2 drops/well). Cells were imaged using a Zeiss LSM 700 laser-scanning confocal fluorescence microscope.

### BioID

Cells were cultured for a total of 48h on five or ten 15 cm plates (for inducible or endogenous BirA* lines, respectively) in 5% FBS medium and were treated with 50 µM Biotin during the final 24h. Cell pellets were lysed in 1 mL/plate ice-cold RIPA lysis buffer (50 mM Tris-HCl pH 7.5, 150 mM NaCl, 1% NP-40, 1 mM EDTA, 1 mM EGTA, 0.1% SDS, protease inhibitor cocktail (P8340) 1:500, and 0.5% Sodium deoxycholate, supplemented with 250U of benzonase) for 1h at 4ºC. The lysates were sonicated at 4ºC using three 30s bursts, with 1’ pauses, at 30% amplitude. The lysates were then centrifuged at 4ºC for 30’ at 20k × g. Streptavidin sepharose beads (17–5113–01, GE) were washed 3x with 1 mL RIPA buffer and pelleted at 2000 × g for 2’. Biotinylated proteins were captured from cleared lysates by incubating 3h at 4ºC, with 10 µL of packed beads per 15 cm plate. The beads were pelleted at 2000 × g for 2’ and transferred to 1.5 mL tubes in 1 mL RIPA buffer. 6 washes were performed with 1 mL 50 mM ammonium bicarbonate. Beads were then resuspended in 200 l ammonium bicarbonate and 1 µg of trypsin (PRV5111, Promega) was added. The samples were incubated at 37ºC overnight, followed by an additional 2–4h incubation with an additional 0.5 µg trypsin. The beads were pelleted, and the supernatant was transferred to a new tube. The beads were then rinsed twice with 150 µl ammonium bicarbonate and the wash fractions were pooled together with the supernatant. The samples were lyophilized in a speed-vac, resuspended in 8 µl formic acid, and 1/4 was used per mass spectrometric analysis.

### Mass Spectrometry

All BioID samples and controls were analyzed by mass spectrometry with at least three biological replicates. Liquid chromatography was conducted using a home-made trap-column (5 cm × 200 µm inner diameter) and a home-made analytical column (50 cm × 50 µm inner diameter; Monitor 5µm 100A C18 resin). A 2h reversed-phase gradient was run at 70 nl/min on a Thermo Fisher Ultimate 3000 RSLC Nano UPLC system coupled to a Thermo QExactive HF quadrupole-Orbitrap mass spectrometer. A parent ion scan was performed using a resolving power of 120 000, and up to the 10 most intense peaks were selected for MS/MS (minimum ion count of 1000 for activation) using higher energy collision induced dissociation (HCD) fragmentation. Dynamic exclusion was activated such that MS/MS of the same m/z (within a range of 10 ppm; exclusion list size = 500) detected twice within 5s were excluded from analysis for 30s.

### Mass Spectrometric Data Analysis

Mass spectrometric data from the Thermo QExactive HF quadrupole-Orbitrap were stored, searched and analyzed using the ProHits laboratory information management system (46). Raw files were searched using Mascot and Comet, against the MouseV53cRapRevTagA database, which includes common contaminants (47, 48). The database parameters were set to search for tryptic cleavages, allowing up to 1 missed cleavage site per peptide, with a parent MS tolerance of 12 ppm for precursors with charges of 2+ to 4+ and a fragment ion tolerance of ±0.15 amu. Variable modifications were selected for deamidated asparagine/glutamine and oxidized methionine. The results from each search were statistically validated through the Trans-Proteomic Pipeline, using iProphet (49, 50). SAINTexpress was used to calculate the probability of each potential proximal–protein from background contaminants using default parameters (51). Controls were compressed to 2 samples. 2 unique peptides and a minimum iProphet probability of 0.95 were required for protein identification. SAINTexpress data were analyzed and visualized using ProHits-Viz (52).

## Author Contributions

BWD and SM designed the experiments, conducted experiments and wrote the paper. CS, VG, EP, VF and DN conducted experiments. A-CG., BL, BR, and CJW provided BioID bioinformatic analyses. YL, SX and RW performed mass spectro-metric data acquisition.

## Acknowledgements

Funding for this study and required infrastructure was provided by the Canadian Institutes of Health Research to BWD (MOP133610), the Canada Research Chairs Program (BWD), the Ontario Ministry of Research and Innovation (BWD), the Canada Foundation for Innovation (BWD), and the OCRiT Project: Ontario Ministry of Economic Development and Innovation (BWD). Bioinformatics analysis was performed at the Network Biology Collaborative Centre at the Lunenfeld-Tanenbaum Research Institute, a facility supported by Canada Foundation for Innovation funding, by the Ontarian Government and by Genome Canada and Ontario Genomics (OGI-097, OGI-139).

## References

1. Hans Clevers and Roel Nusse. Wnt/*β*–Catenin Signaling and Disease. Cell, 149(6):1192–1205, jun 2012.

2. Roel Nusse and Hans Clevers. Wnt/*β*–Catenin Signaling, Disease, and Emerging Therapeutic Modalities, 2017.

3. Ken M Cadigan and Marian L Waterman. TCF/LEFs and Wnt Signaling in the Nucleus. Cold Spring Harb Perspect Biol, 4(11):a007906–a007906, 2012.

4. Britta Wallmen, Monika Schrempp, and Andreas Hecht. Intrinsic properties of Tcf1 and Tcf4 splice variants determine cell-type-specific Wnt/*β*–catenin target gene expression. Nucleic Acids Research, 40(19):9455–9469, 2012.

5. Steven Moreira, Enio Polena, Victor Gordon, Solen Abdulla, Sujeivan Mahendram, Jiayi Cao, Alexandre Blais, Geoffrey A G.A. Wood, Anna Dvorkin-Gheva, and Bradley W B.W. Bradley W Doble. A Single TCF Transcription Factor, Regardless of Its Activation Capacity, Is Sufficient for Effective Trilineage Differentiation of ESCs. Cell Reports, 20(10):2424–2438, 2017.

6. Laura Pereira, Fei Yi, and Bradley J Merrill. Repression of Nanog gene transcription by Tcf3 limits embryonic stem cell self-renewal. Molecular and Cellular Biology, 26(20):7479–7491, 2006.

7. Megan F Cole, Sarah E Johnstone, Jamie J Newman, Michael H Kagey, and Richard A Young. Tcf3 is an integral component of the core regulatory circuitry of embryonic stem cells. Genes & Development, 22(6):746–755, 2008.

8. Alexander Marson, Stuart S Levine, Megan F Cole, Garrett M Frampton, Tobias Brambrink, Sarah Johnstone, Matthew G Guenther, Wendy K Johnston, Marius Wernig, Jamie Newman, J Mauro Calabrese, Lucas M Dennis, Thomas L Volkert, Sumeet Gupta, Jennifer Love, Nancy Hannett, Phillip A Sharp, David P Bartel, Rudolf Jaenisch, and Richard A Young. Connecting microRNA Genes to the Core Transcriptional Regulatory Circuitry of Embryonic Stem Cells. Cell, 134(3):521–533, 2008.

9. J Wray, T Kalkan, S Gomez-Lopez, D Eckardt, A Cook, R Kemler, and A Smith. Inhibition of glycogen synthase kinase-3 alleviates Tcf3 repression of the pluripotency network and increases embryonic stem cell resistance to differentiation. Nat Cell Biol, 13(7):838–845, 2011.

10. Fei Yi, Laura Pereira, Jackson A Hoffman, Brian R Shy, Courtney M Yuen, David R Liu, and Bradley J Merrill. Opposing effects of Tcf3 and Tcf1 control Wnt stimulation of embryonic stem cell self-renewal. Nat Cell Biol, 13(7):762–770, 2011.

11. Brian R Shy, Chun-I I Wu, Galina F Khramtsova, Jenny Y Zhang, Olufunmilayo I Olopade, Kathleen H Goss, and Bradley J Merrill. Regulation of Tcf7l1 DNA Binding and Protein Stability as Principal Mechanisms of Wnt/*β*–Catenin Signaling. CellReports, 4(1):1–9, 2013.

12. Yaser Atlasi, Rubina Noori, Claudia Gaspar, Patrick Franken, Andrea Sacchetti, Haleh Rafati, Tokameh Mahmoudi, Charles Decraene, George A Calin, Bradley J Merrill, and Riccardo Fodde. Wnt Signaling Regulates the Lineage Differentiation Potential of Mouse Embryonic Stem Cells through Tcf3 Down-Regulation. PLoS Genetics, 9(5):e1003424, 2013.

13. Mark Brannon, Jeffrey D Brown, Rebecca Bates, David Kimelman, and Randall T Moon. XCtBP is a XTcf-3 co-repressor with roles throughout Xenopus development. Development (Cambridge, England), 126(14):3159–3170, 1999.

14. Melanie A Eshelman, Meera Shah, Wesley M Raup-Konsavage, Sherri A Rennoll, and Gregory S Yochum. TCF7L1 recruits CtBP and HDAC1 to repress DICKKOPF4 gene expression in human colorectal cancer cells. Biochemical and Biophysical Research Communications, 487(3):716–722, 2017.

15. Wen-Hui Lien, Lisa Polak, Mingyan Lin, Kenneth Lay, Deyou Zheng, and Elaine Fuchs. In vivo transcriptional governance of hair follicle stem cells by canonical Wnt regulators. Nature Cell Biology, 16(2):179–190, 2014.

16. Wai-Leong Tam, Chin Yan Lim, Jianyong Han, Jinqiu Zhang, Yen-Sin Ang, Huck-Hui Ng, Henry Yang, and Bing Lim. Tcf3 Regulates Embryonic Stem Cell Pluripotency and Self-Renewal by the Transcriptional Control of Multiple Lineage Pathways. Stem cells (Dayton, Ohio), 26(8):2019–2031, 2008.

17. Chun-I Wu, Jackson a Hoffman, Brian R Shy, Erin M Ford, Elaine Fuchs, Hoang Nguyen, and Bradley J Merrill. Function of Wnt/*β*–catenin in counteracting Tcf3 repression through the Tcf3-*β*–catenin interaction. Development (Cambridge, England), 139(12):2118–2129, 2012.

18. N Barker, a Hurlstone, H Musisi, a Miles, M Bienz, and H Clevers. The chromatin remodelling factor Brg-1 interacts with beta-catenin to promote target gene activation. The EMBO Journal, 20(17):4935–4943, 2001.

19. a Hecht. The p300/CBP acetyltransferases function as transcriptional coactivators of beta-catenin in vertebrates. The EMBO Journal, 19(8):1839–1850, 2000.

20. Thomas Kramps, Oliver Peter, Erich Brunner, Denise Nellen, Barbara Froesch, Sandipan Chatterjee, Maximilien Murone, Stephanie Züllig, and Konrad Basler. Wnt/Wingless signal-ing requires BCL9/legless-mediated recruitment of pygopus to the nuclear *β*–catenin-TCF complex. Cell, 109(1):47–60, 2002.

21. Ken Ichi Takemaru and Randall T Moon. The transcriptional coactivator CBP interacts with *β*–catenin to activate gene expression. Journal of Cell Biology, 149(2):249–254, 2000.

22. Barry Thompson, Fiona Townsley, Rina Rosin-Arbesfeld, Hannah Musisi, and Mariann Bienz. A new nuclear component of the Wnt signalling pathway. Nature Cell Biology, 4(5): 367–373, 2002.

23. Kyle J Roux, Dae In Kim, Manfred Raida, and Brian Burke. A promiscuous biotin ligase fusion protein identifies proximal and interacting proteins in mammalian cells. The Journal of Cell Biology, 196(6):801–810, 2012.

24. Dae In Kim, B. KC, Wenhong Zhu, Khatereh Motamedchaboki, Valérie Doye, and Kyle J Roux. Probing nuclear pore complex architecture with proximity-dependent biotinylation. Proceedings of the National Academy of Sciences, 111(24):E2453–E2461, 2014.

25. Fei Yi, Laura Pereira, and Bradley James Merrill. Tcf3 Functions as a Steady-State Limiter of Transcriptional Programs of Mouse Embryonic Stem Cell Self-Renewal. Stem Cells, 26 (8):1951–1960, 2008.

26. Danette L Daniels and William I Weis. Beta-catenin directly displaces Groucho/TLE repressors from Tcf/Lef in Wnt-mediated transcription activation. Nature Publishing Group, 12(4): 364–371, 2005.

27. Jayanth V Chodaparambil, Kira T Pate, Margretta R.D. Hepler, Becky P Tsai, Uma M Muthurajan, Karolin Luger, Marian L Waterman, and William I Weis. Molecular functions of the TLE tetramerization domain in Wnt target gene repression. EMBO Journal, 33(7):719–731, 2014.

28. Laurens M van Tienen, Juliusz Mieszczanek, Marc Fiedler, Trevor J Rutherford, and Mariann Bienz. Constitutive scaffolding of multiple Wnt enhanceosome components by legless/BCL9. eLife, 6:1–23, 2017.

29. Jackson a Hoffman, Chun-I Wu, and Bradley J Merrill. Tcf7l1 prepares epiblast cells in the gastrulating mouse embryo for lineage specification. Development (Cambridge, England), 140(8):1665–1675, 2013.

30. Miguel Fidalgo, Xin Huang, Diana Guallar, Carlos Sanchez-Priego, Victor Julian Valdes, Arven Saunders, Junjun Ding, Wen Shu Wu, Carlos Clavel, and Jianlong Wang. Zfp281 Coordinates Opposing Functions of Tet1 and Tet2 in Pluripotent States. Cell Stem Cell, 19 (3):355–369, 2016.

31. Feng Xiao, Bing Liao, Jing Hu, Shuang Li, Haixin Zhao, Ming Sun, Junjie Gu, and Ying Jin. JMJD1C Ensures Mouse Embryonic Stem Cell Self-Renewal and Somatic Cell Reprogramming through Controlling MicroRNA Expression. Stem Cell Reports, 9(3):927–942, sep 2017.

32. Jianchang Yang, Li Chai, Taylor C Fowles, Zaida Alipio, Dan Xu, Louis M Fink, David C Ward, and Yupo Ma. Genome-wide analysis reveals Sall4 to be a major regulator of pluripotency in murine-embryonic stem cells. Proceedings of the National Academy of Sciences of the United States of America, 105(50):19756–19761, 2008.

33. J Q Zhang, W L Tam, G Q Tong, Q Wu, H Y Chan, B S Soh, Y F Lou, J C Yang, Y P Ma, L Chai, H H Ng, T Lufkin, P Robson, and B Lim. Sall4 modulates embryonic stem cell pluripotency and early embryonic development by the transcriptional regulation of Pou5f1. Nature Cell Biology, 8(10):1114–U125, 2006.

34. Sridhar Rao, S Zhen, Sergei Roumiantsev, Lindsay T McDonald, Guo-Cheng C Yuan, Stuart H Orkin, Zhen Shao, Sergei Roumiantsev, Lindsay T McDonald, Guo-Cheng C Yuan, and Stuart H Orkin. Differential Roles of Sall4 Isoforms in Embryonic Stem Cell Pluripotency. Molecular and Cellular Biology, 30(22):5364–5380, 2010.

35. Johann Böhm, Claudio Sustmann, Christian Wilhelm, and Jürgen Kohlhase. SALL4 is directly activated by TCF/LEF in the canonical Wnt signaling pathway. Biochemical and Biophysical Research Communications, 348(3):898–907, 2006.

36. Yupo Ma, Wei Cui, Jianchang Yang, Jun Qu, Chunhui Di, Hesham M. Amin, Raymond Lai, Jerome Ritz, Diane S. Krause, and Li Chai. SALL4, a novel oncogene, is constitutively expressed in human acute myeloid leukemia (AML) and induces AML in transgenic mice. Blood, 108(8):2726–2735, 2006.

37. R T Collins and J E Treisman. Osa-containing Brahma chromatin remodeling complexes are required for the repression of Wingless target genes. Genes and Development, 14(24): 3140–3152, 2000.

38. Aliaksei Z Holik, Madeleine Young, Joanna Krzystyniak, Geraint T Williams, Daniel Metzger, Boris Y Shorning, and Alan R Clarke. Brg1 Loss Attenuates Aberrant Wnt-Signalling and Prevents Wnt-Dependent Tumourigenesis in the Murine Small Intestine. PLoS Genetics, 10 (7), 2014.

39. Christoph Schaniel, Yen-Sin Ang, Kajan Ratnakumar, Catherine Cormier, Taneisha James, Emily Bernstein, Ihor R Lemischka, and Patrick J Paddison. Smarcc1/Baf155 Couples Self-Renewal Gene Repression with Changes in Chromatin Structure in Mouse Embryonic Stem Cells. Stem Cells, 27(12):N/A–N/A, 2009.

40. L Ho, J L Ronan, J Wu, B T Staahl, L Chen, A Kuo, J Lessard, A I Nesvizhskii, J Ranish, and G R Crabtree. An embryonic stem cell chromatin remodeling complex, esBAF, is essential for embryonic stem cell self-renewal and pluripotency. Proceedings of the National Academy of Sciences, 106(13):5181–5186, 2009.

41. Gillian Morrison, Roberta Scognamiglio, Andreas Trumpp, and Austin Smith. Convergence of cMyc and -catenin on Tcf7l1 enables endoderm specification. The EMBO Journal, 35(3): 356–368, 2016.

42. S Ogawa, J Lozach, K Jepsen, D Sawka-Verhelle, V Perissi, R Sasik, D W Rose, R S Johnson, M G Rosenfeld, and C K Glass. A nuclear receptor corepressor transcriptional checkpoint controlling activator protein 1-dependent gene networks required for macrophage activation. Proceedings of the National Academy of Sciences, 101(40):14461–14466, 2004.

43. Valentina Perissi, Aneel Aggarwal, Christopher K Glass, David W Rose, and Michael G Rosenfeld. A Corepressor/Coactivator Exchange Complex Required for Transcriptional Activation by Nuclear Receptors and Other Regulated Transcription Factors. Cell, 116(4): 511–526, 2004.

44. Valentina Perissi, Claudio Scafoglio, Jie Zhang, Kenneth A Ohgi, David W Rose, Christopher K Glass, and Michael G Rosenfeld. TBL1 and TBLR1 Phosphorylation on Regulated Gene Promoters Overcomes Dual CtBP and NCoR/SMRT Transcriptional Repression Checkpoints. Molecular Cell, 29(6):755–766, 2008.

45. Jiong Li and Cun Yu Wang. TBL1-TBLR1 and *β*–catenin recruit each other to Wnt target-gene promoter for transcription activation and oncogenesis. Nature Cell Biology, 10(2): 160–169, 2008.

46. Guomin Liu, Jianping Zhang, Brett Larsen, Chris Stark, Ashton Breitkreutz, Zhen-Yuan Lin, Bobby-Joe Breitkreutz, Yongmei Ding, Karen Colwill, Adrian Pasculescu, Tony Pawson, Jeffrey L Wrana, Alexey I Nesvizhskii, Brian Raught, Mike Tyers, and Anne-Claude Gingras. ProHits: integrated software for mass spectrometry-based interaction proteomics. Nature Biotechnology, 28(10):1015–1017, 2010.

47. Jimmy K Eng, Tahmina A Jahan, and Michael R Hoopmann. Comet: An open-source MS/MS sequence database search tool. Proteomics, 13(1):22–24, 2013.

48. David N Perkins, Darryl J.C. Pappin, David M Creasy, and John S Cottrell. Probability-based protein identification by searching sequence databases using mass spectrometry data. In Electrophoresis, volume 20, pages 3551–3567, 1999.

49. Eric W Deutsch, Luis Mendoza, David Shteynberg, Terry Farrah, Henry Lam, Natalie Tasman, Zhi Sun, Erik Nilsson, Brian Pratt, Bryan Prazen, Jimmy K Eng, Daniel B Martin, Alexey I Nesvizhskii, and Ruedi Aebersold. A guided tour of the Trans-Proteomic Pipeline, 2010.

50. David Shteynberg, Eric W Deutsch, Henry Lam, Jimmy K Eng, Zhi Sun, Natalie Tasman, Luis Mendoza, Robert L Moritz, Ruedi Aebersold, and Alexey I Nesvizhskii. iProphet: Multilevel Integrative Analysis of Shotgun Proteomic Data Improves Peptide and Protein Identification Rates and Error Estimates. Molecular & Cellular Proteomics, 10(12):M111.007690, 2011.

51. Hyungwon Choi, Brett Larsen, Zhen-Yuan Lin, Ashton Breitkreutz, Dattatreya Mellacheruvu, Damian Fermin, Zhaohui S Qin, Mike Tyers, Anne-Claude Gingras, and Alexey I Nesvizhskii. SAINT: probabilistic scoring of affinity purification–mass spectrometry data. Nature Methods, 8(1):70–73, 2010.

52. James D.R. Knight, Hyungwon Choi, Gagan D Gupta, Laurence Pelletier, Brian Raught, Alexey I Nesvizhskii, and Anne Claude Gingras. ProHits-viz: A suite of web tools for visualizing interaction proteomics data, 2017.

